# Structural and functional alterations of neuromuscular synapses in a mouse model of *ACTA1* congenital myopathy

**DOI:** 10.1101/2022.02.23.481672

**Authors:** Yun Liu, Weichun Lin

**Affiliations:** Department of Neuroscience, UT Southwestern Medical Center, 6000 Harry Hines Blvd, Dallas, TX 75390-9111, USA

## Abstract

Mutations in skeletal muscle α-actin (Acta1) cause a variety of myopathies. In a mouse model of congenital myopathy, heterozygous Acta1 (H40Y) knock-in (*Acta1*^+*/Ki*^) mice exhibit features of human nemaline myopathy, including premature lethality, severe muscle weakness, reduced mobility, and the presence of nemaline rods in muscle fibers. In this study, we investigated the structure and function of the neuromuscular junction (NMJ) in the *Acta1*^+*/Ki*^ mice. We found marked impairments in NMJ structure in the mutant mice, including fragmented endplates and nerve terminals, reduced density of acetylcholine receptors (AChRs) on endplate membranes, reduced nerve occupancy at endplates, and increased numbers of muscle fiber subsynaptic nuclei. We compared the NMJs in three different types of muscles – the extensor digitorum longus (EDL, composed of fast-twitch muscle fibers), soleus (Sol, enriched in slow-twitch fibers) and the triangularis sterni muscle (TS, a mixed fiber type muscle). Among these three types of muscles, EDL was affected to the greatest extent, suggesting that fast-twitch fibers may be most susceptible to NMJ fragmentation in *Acta1*^+*/Ki*^ nemaline myopathy.

Electrophysiological analysis of mutant NMJs showed a reduced quantal size (reduced mEPP amplitude), increased mEPP frequency, and increased quantal content, but normal EPP amplitude compared to wild type (*WT*) NMJs. The results suggest that affected synapses may have undergone homeostatic compensation to maintain normal levels of neurotransmitter release. In addition, paired-pulse facilitation was reduced and synaptic depression under repetitive nerve stimulation was enhanced, indicating shortterm synaptic plasticity was compromised in the mutant mice.

**Key points:**

- Mice heterozygous for an Acta1 (H40Y) knock-in mutation exhibit clinical features of human nemaline myopathy. We report structural and functional alterations of neuromuscular synapses in these mutant mice. The NMJ impairments include endplate fragmentation, reduced endplate nerve occupancy, and increased numbers of subsynaptic nuclei in muscle fibers.
- Neuromuscular synaptic transmission was compromised – demonstrating both increased quantal content and changes in short-term synaptic plasticity.
- Increases in spontaneous neurotransmitter release and quantal content suggest homeostatic compensation of synapses to maintain normal transmitter release in the mutant NMJs.

## Introduction

Mammalian actin is composed of six isoforms, including four muscle actins [(α_skeletal_-actin (Acta1), α_cardiac_-actin (Actc1), α_smooth_-actin (Acta2), and γ_smooth_-actin (Actg2)] and two non-muscle actins [β_cyto_-actin (Actb) and γ_cyto_-actin (Actg1)] (Perrin & Ervasti, 2010; Kashina, 2020). Skeletal muscle α-actin is the principle component of the thin filaments in adult skeletal muscle. The interaction between skeletal α-actin in thin filaments and myosin in thick filaments produces force, leading to muscle contractions. Mutant mice lacking skeletal α-actin (*Acta1*^-/-^) display significantly reduced muscle force production and die during the early neonatal period (Crawford *et al*., 2002). Clinically, mutations in the skeletal *Acta1* gene are associated with a variety of congenital myopathies. These include nemaline myopathy, intranuclear rod myopathy, actin-accumulation myopathy, central core disease and congenital fiber type disproportion (Nowak *et al*., 1999; Ilkovski *et al*., 2001; Sparrow *et al*., 2003; Agrawal *et al*., 2004; Laing *et al*., 2004; Wallefeld *et al*., 2006; Nowak *et al*., 2013; Yang *et al*., 2016). The lethal human missense mutation H40Y, located in the DNase I-binding loop of the actin sub-domain 2, greatly disrupts the binding of actin filaments to myosin molecules and therefore leads to contractile dysfunction and severe muscle weakness (Chan *et al*., 2016). Human patients with the *ACTA1* (H40Y) mutation develop severe muscle weakness and nemaline myopathies (Nowak *et al*., 1999). In a mouse model of *ACTA1* nemaline myopathy, mice heterozygous for the *Acta1* (H40Y) knock-in mutation (*Acta1*^+*/Ki*^) exhibit clinical pathological characteristic of patients with this mutation, including premature lethality, severe muscle weakness, reduced mobility, the presence of nemaline rods, and muscle fiber atrophy (Nguyen *et al*., 2011).

Muscle pathology, as exhibited in *Acta1*^+*/Ki*^ mutant mice (Nguyen *et al*., 2011; Lindqvist *et al*., 2013), may impact the structure and function of the neuromuscular junction (NMJ), which is the synaptic connection between a motor neuron and a skeletal muscle fiber. Indeed, alterations in the NMJ have been widely studied in physiological processes such as aging (Jang & Van Remmen, 2011; Gonzalez-Freire *et al*., 2014; Rudolf *et al*., 2014; Li *et al*., 2018) and in various neuromuscular disorders such as Duchenne muscular dystrophy (DMD) (Theroux *et al*., 2008; van der Pijl *et al*., 2016; Lovering *et al*., 2020). In this study, we sought to assess NMJ morphology and function in *Acta1*^+*/Ki*^ mutant mice. We found significant changes in the structure of the NMJ when compared to *WT*. The alterations include fragmented endplates, fragmented nerve terminals, loss of acetylcholine receptors (AChR), reduced nerve occupancy at the endplate, and increased numbers of subsynaptic nuclei. Furthermore, neuromuscular synaptic transmission was also markedly altered in *Acta1*^+*/Ki*^ mutant mice. The size of spontaneous synaptic transmission, as measured by miniature end-plate potential (mEPP) amplitude, was reduced, while the mEPP frequency was increased. The combined effect of the reduction in mEPP size and increase in quantal content yielded normal amplitude for the evoked endplate potential (EPP). In addition, synaptic plasticity was compromised in *Acta1*^+*/Ki*^ mutant mice, evidenced by a reduction in pair-pulse facilitation and an increase in synaptic depression in response to trains of nerve stimuli. Together, these findings document profound changes in both the structure and function of the NMJs in *Acta1*^+*/Ki*^ mutant mice.

## Methods and Materials

### Mice

Heterozygous *Acta1* (H40Y) mutant mice (also known as Acta1^tm1(H40Y;neo)Hrd^, hereafter as *Acta1*^+*/Ki*^) were obtained from the Jackson Laboratory at Bar Harbor, Main, USA (strain # 018284, MGI: 5424775). These mice were originally generated in the laboratory of Dr. Edna Hardeman (The University of New South Wales, Sydney, Australia) (Nguyen *et al*., 2011). In these mutant mice, the endogenous ACTA1 was replaced by a mutant ACTA1 [Acta1 (H40Y) knock-in allele]) having a single amino acid substitution of histidine to tyrosine at codon 40 (H40Y). *Acta1*^+*/Ki*^ mice exhibited premature lethality (Nguyen *et al*., 2011). These mice were bred with C57BL/6J mice to generate *Acta1*^+*/Ki*^ and littermate wild type (*Acta1*^+/+^, *WT*) controls for experiments. All experimental protocols followed National Institutes of Health Guidelines and were approved by the University of Texas Southwestern Institutional Animal Care and Use Committee.

### Immunofluorescence staining

Whole mount immunofluorescence staining was carried out as previously described (Liu *et al*., 2008). Soleus (Sol), extensor digitorum longus (EDL), and triangularis sterni (TS) muscles from *Acta1*^+*/Ki*^ and wildtype littermate mice were fixed with 2% paraformaldehyde in 0.1 M phosphate buffer (pH 7.3) overnight at 4°C. Muscle samples were extensively washed with PBS and then incubated with Texas Red-conjugated α-bungarotoxin (α-bgt) (2 nM, Invitrogen, Carlsbad, California, USA) for 30 minutes at room temperature. Samples were then incubated with primary antibodies overnight at 4°C. The following polyclonal antibodies were used: anti-syntaxin 1 (I375) and anti-synaptotagmin 2 (I735) (generous gifts from Dr. Thomas Südhof, Stanford University School of Medicine, Palo Alto, CA, USA), and anti-AChE (generous gifts from Dr. Palmer Taylor, Skaggs School of Pharmacy & Pharmaceutical Sciences, UC San Diego, CA, USA). All primary antibodies were diluted by 1:1000 in antibody dilution buffer (500 mM NaCl, 0.01 M phosphate buffer, 3% BSA, and 0.01% thimerosal). After extensive washes, muscle samples were then incubated with fluorescein isothiocyanate (FITC)-conjugated goat anti-rabbit IgG (1:600, Jackson ImmunoResearch Laboratories, Inc., West Grove, PA, USA) overnight at 4°C. For the nuclei labeling experiment, muscle samples were further incubated in ToPro-3 (1:3000, Eugene, Oregon, USA) for 30 minutes at room temperature. Muscle samples were mounted in the Vectashield mounting medium (H-1000, Vector Laboratories, Inc., Burlingame, CA, USA). Images were captured using a Zeiss LSM 880 confocal microscope.

### Morphometric analysis

Confocal images captured at high magnification (63×/1.4 Oil DIC M27) were used for the following morphometric analyses. The endplate area was measured as the thresholded area of AChR labeled with α-bungarotoxin. The synaptic area was measured as the polygon shaped region that was manually traced around the thresholded signal of AChR labeled with α-bungarotoxin. A dispersion index was obtained by dividing the synaptic area by the endplate area. To calculate nerve occupancy of a NMJ, the ratio of presynaptic (nerve) area to postsynaptic (endplate) area was calculated (Pratt *et al*., 2015). Presynaptic area was measured as the thresholded area of the nerve labeled with antibody against synaptotagmin 2. To analyze the nerve complexity within a NMJ, we used antibody against syntaxin 1 to label the nerves. A line was drawn along the longest axis across the NMJ and the number of intersections between the line and the axon labeled by syntaxin 1 was counted (Liu *et al*., 2019).

### Electrophysiology

Electrophysiological analyses were performed on *Acta1*^+*/Ki*^ and *WT* littermates at 2 months of age. Intracellular recordings were carried out at room temperature as previously described (Liu *et al*., 2011). Briefly, the samples of tibial nerve-Sol muscle and fibular nerve-EDL muscle were dissected and mounted on a Sylgard coated dish, and bathed in oxygenated (95% O_2_, 5% CO_2_) Ringer’s solution (136.8 mM NaCl, 5 mM KCl, 12 mM NaHCO_3_, 1 mM NaH_2_PO_4_, 1 mM MgCl_2_, 2 mM CaCl_2_, and 11 mM d-glucose, pH 7.3). Under a water-immersion objective (Olympus BX51WI), end-plate regions were identified and impaled with glass microelectrodes (resistance 20 - 40 MΩ) filled with 2 M potassium citrate and 10 mM potassium chloride. Supra threshold stimuli (2 - 5 V, 0.1ms) were applied to the nerve via a glass suction electrode connected to an extracellular stimulator (SD9, Grass-Telefactor, West Warwick, RI). To prevent muscle contractions, μ-conotoxin GIIIB (2 μM; Peptides International) was added to the bath solution 30 minutes prior to recording. Miniature endplate potentials (mEPPs) and evoked endplate potentials (EPPs) were acquired with an intracellular amplifier (AxoClamp-2B) and digitized with Digidata 1332A (Molecular Devices, Sunnyvale, CA, USA). Data were analyzed with pClamp 10.7 (Molecular Devices) and Mini Analysis Program (Synaptosoft, Inc., Decatur, GA). Quantal content (the number of acetylcholine quanta released in response to a single nerve impulse) was estimated by dividing the mean amplitude of EPPs by the mean amplitude of mEPPs of the same cell (Boyd & Martin, 1956; Wood & Slater, 2001). Rise time of mEPPs or EPPs was calculated as the time taken by the membrane potential to rise from 10% to 90% of the peak value of mEPPs or EPPs. Decay time of mEPPs was calculated as the time taken by the membrane potential to decay from 100% to 50% of the peak value of mEPPs. Decay time of EPPs was calculated as the time taken by the membrane potential to decay from 90% to 10% of the peak value of EPPs.

### Data analyses

Data were presented as mean ± standard error of the mean (SEM). For quantitative morphometric analyses, **t**he percentage of fragmented NMJs, average fragment number per endplate, endplate size, dispersion index of AChR, percentage of the NMJ with faint or loss of AChR stain, nerve occupancy to the endplate, nerve intersection number and subsynaptic nuclei density of Sol, EDL and TS muscles were compared between wildtype (*Acta1*^+/+^) and mutant (*Acta1*^+*/Ki*^) mice by using student *t*-test. For spontaneous and evoked neurotransmission analyses, the mEPP frequency, amplitude, rise time (10%~90%), decay time (100%~50%) and EPP amplitude, quantal content, EPP rise time (10%~90%) and decay time (90%~10%) of Sol and EDL muscles were compared by using student *t*-test. For paired-pulse facilitation analyses, the ratios of the second to the first EPP amplitude (EPP(2)/EPP(1)) of soleus and EDL muscles were compared by using student paired *t*-test. For synaptic short-term depression analyses, the ratios of EPP(n) to EPP(1) of soleus and EDL were compared by using student paired *t*-test. The difference between wildtype and mutant mice was considered statistically significant if the P-value was less than 0.05.

## Results

### Endplates become fragmented in *Acta1*^+*/Ki*^ mice

Previous studies have shown that *Acta1*^+*/Ki*^ mice exhibit clinical features resembling human nemaline myopathies, including premature death, severe muscle weakness, reduced mobility, nemaline rods in skeletal muscle fibers, and muscle regeneration (Nguyen *et al*., 2011). We set out to examine the structure and function of the NMJ in *Acta1*^+*/Ki*^ mice. Due to the early mortality of *Acta1*^+*/Ki*^ mice, we performed our initial experiments in mice of 2 months of age.

First, we carried out morphological analyses of the NMJs in soleus (Sol), extensor digitorum longus (EDL), and triangularis sterni (TS) muscles. In *WT* muscles, endplates typically appeared continuous and pretzel-shaped (top row in Fig. 1A). In contrast, endplates in EDL and TS muscles of *Acta1*^+*/Ki*^ mice appeared fragmented (arrowheads in Fig. 1A). In contrast, the endplates in Sol muscles appeared unaffected in *Acta1*^+*/Ki*^ mice. Quantitative analyses showed that 74.5% of endplates in EDL and 56.3% in TS were fragmented in *Acta1*^+*/Ki*^ mice (Fig. 1B). However, only 2.8% of endplates in the Sol muscles of *Acta1*^+*/Ki*^ mice were fragmented (Fig. 1B). To better quantify the degree of endplate fragmentation, we counted the average fragment number per endplate. On average, we found 1 fragment per endplate in Sol muscles in both *WT* and *Acta1*^+*/Ki*^ mice. However, the fragmented endplate numbers were increased by ~ 7-fold in *Acta1*^+*/Ki*^ EDL and ~ 6-fold in *Acta1*^+*/Ki*^ TS muscles compared with the same muscles in *WT* mice (Fig. 1C). The endplate size was significantly decreased in EDL muscles of *Acta1*^+*/Ki*^ mice compared to in EDL of *WT* mice (Fig. 1D). The pattern of AChRs, labelled by *α*-bungarotoxin, appeared more dispersed in the EDL and TS muscles of *Acta1*^+*/Ki*^ mice, compared to *WT* (Fig 1 E). AChR staining appeared homogenous in *WT* muscles (top panel in Fig. 1F), but heterogeneous in *Acta1*^+*/Ki*^ muscles – some regions were faint or even lacked *α*-bungarotoxin staining (arrowheads in Fig. 1F). This was quantitated in Fig. 1 G: 17%-50% of the NMJ in *Acta1*^+*/Ki*^ muscles were faint or lacked AChR staining, which was significantly more than in *WT* muscles (3%-7%) (Fig 1G). These results suggest a loss of AChRs in *Acta1*^+*/Ki*^ muscles.

**Figure 1.**
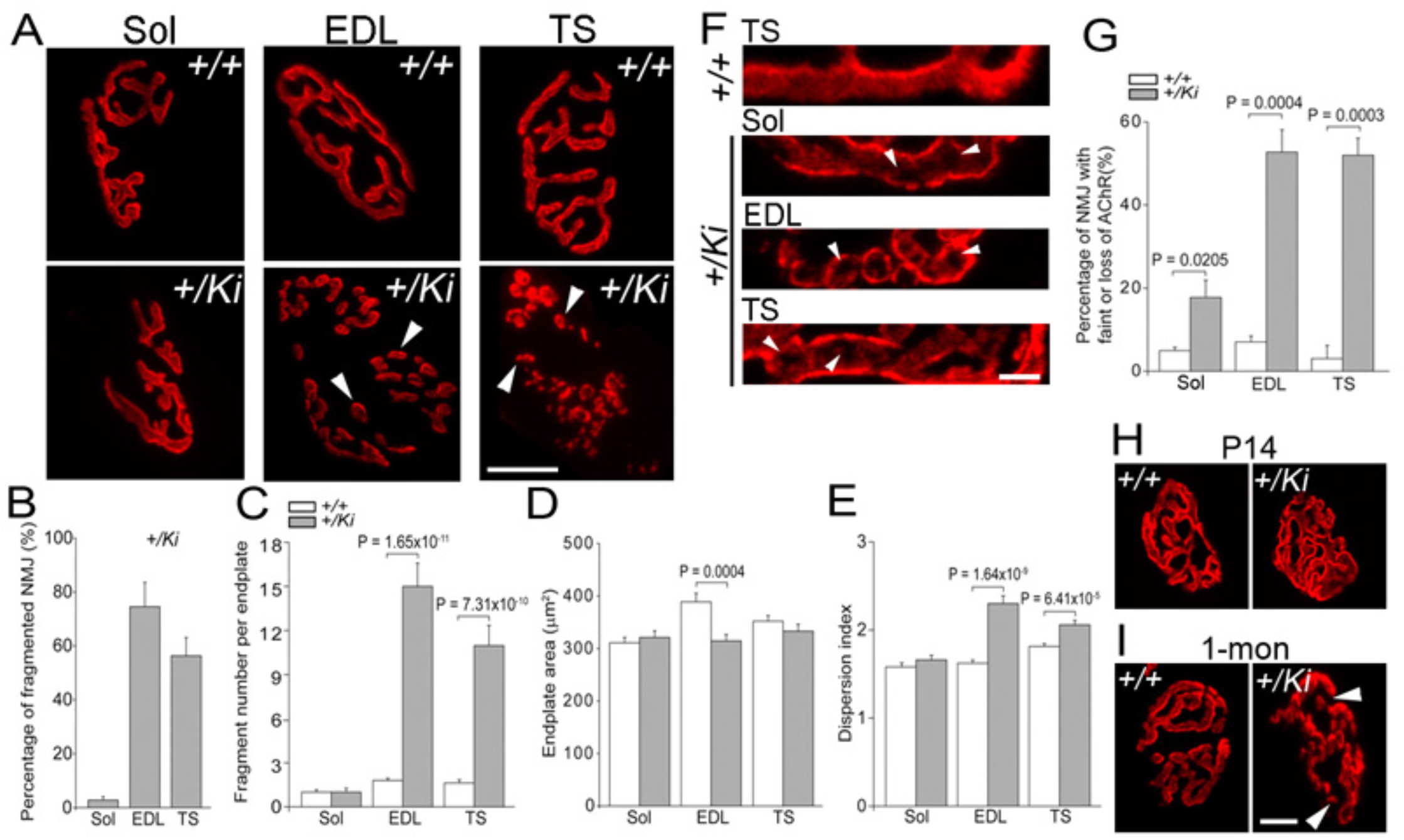
Fragmentation of endplates *in Acta1*^+*/Ki*^ mice. A: Whole mounts of soleus (Sol), EDL and triangularis sterni (TS) muscles of *Acta1*^+*/Ki*^ and *WT* mice (2-month-old) were stained with Texas Red-conjugated *α*-bungarotoxin to label AChRs at the motor endplate. In *WT* muscles, endplates appeared continuous and pretzel shaped. In contrast, in EDL and TS muscles of *Acta1*^+*/Ki*^ mice, endplates were fragmented and contained numerous isolated “islands” of AChRs (arrowheads in A). However, endplates in *ACTα1*^+*/Ki*^ Sol muscle was not notably affected and appeared normal. B: Bar graphs comparing the percentage of fragmented endplates in *Acta1*^+*/Ki*^ mice. Sol: 2.81 ± 1.28 %; EDL: 74.52 ± 9 %; TS: 56.28 ± 6.81 %. An endplate was defined as fragmented when it contained more than 5 fragments. C: A comparison of the average fragment number per endplate between *WT* and *Acta1*^+*/Ki*^ mice. The Sol muscle in *Acta1*^+*/Ki*^ (1 ± 0.26) exhibited similar fragment number per endplate compared with *WT* Sol (1 ± 0.16). In contrast, an average of 15 ± 1.58 and 10.97 ± 1.38 fragments were detected in *Acta1*^+*/Ki*^ EDL and TS, respectively, significantly more than *WT* EDL (1.8 ± 0.15) and TS (1.63 ± 0.21) endplates. D: Measurement of endplate size. The endplate sizes in *Acta1*^+*/Ki*^ Sol (321.17 ± 12.45 μm^2^) and TS (333 ± 13.36 μm^2^) were comparable to those in *WT* Sol (310.94 ± 10.17 μm^2^) and TS (352.08 ± 10.44 μm^2^). However, the endplate size in *Acta1*^+*/Ki*^ EDL (314.47 ± 12.41 μm^2^) was significantly reduced compared with that in *WT* EDL (388.8 ± 16.4 μm^2^). E: The dispersion index of *Acta1*^+*/Ki*^ Sol muscle (1.66 ± 0.05) was comparable to that in *WT* (1.58 ± 0.05). In contrast, the dispersion indices of *Acta1*^+*/Ki*^ EDL (2.3 ± 0.09) and TS (2.06 ± 0.05) muscles were significantly higher than those in *WT* EDL (1.62 ± 0.04) and TS (1.81 ± 0.05). F: Muscles were stained with Texas Red-conjugated *α*-bungarotoxin. The staining was homogenous in *WT* muscles. In contrast, some holes (denoted by arrowheads) in staining lacked AChRs within the endplates in mutant muscles. G: the percentages of the NMJ with faint or loss of AChR staining. The values were significantly higher in *Acta1*^+*/Ki*^ mutant muscles (Sol: 17.76 ± 4.14 %; EDL: 52.72 ± 5.36 %; TS: 51.97 ± 4.11%), compared to those in *WT* muscles (Sol: 4.94 ± 0.82 %; EDL: 7.04 ± 1.41 %; TS: 3.03 ± 3.16 %). H-I: whole mounts of TS muscles from *WT* and *Acta1*^+*/Ki*^ mice at P14 (H) and 1 month (I) were labeled with Texas Red-conjugated *α*-bungarotoxin for AChRs. Endplate fragmentation was detected in mutant muscle at 1 month of age, but not at P14. The numbers of animal (N) and endplate (n) used for quantitative analysis: Sol (*WT*: N = 3, n = 81; *Acta1*^+*/Ki*^: N = 3, n = 94); EDL (*WT*: N = 3, n = 78; *Acta1*^+*/Ki*^: N = 3, n = 151). TS (*WT*: N = 3, n = 173; *Acta1*^+*/Ki*^: N = 3, n = 173). Scale bars: A, 20 μm; F, 5 μm; H and I, 10 μm.

To determine if the fragmentation of endplates occurred prior to 2 months of age, we examined TS muscles of mice at postnatal 14 days (P14) and 1 month old (1-mon). At P14, NMJs undergo postnatal maturation and endplates transform from a “plaque-like” shape to a “pretzel-like” shape (Sanes & Lichtman, 1999; Marques *et al*., 2000). At P14, endplates in both *WT* and *Acta1*^+*/Ki*^ mice exhibited a “pretzel-like” shape (Fig1 H). No fragmentation was observed at P14. At 1 month of age, fragmented endplates were detected in *Acta1*^+*/Ki*^ mice but not in *WT* (Fig. 1I). Together, these results suggest that endplates of *Acta1*^+*/Ki*^ mice developed normally up to P14, but then begin to deteriorate, showing signs of fragmentation by 1 month of age.

Acetylcholinesterase (AChE) is the enzyme that catalyzes hydrolysis of the neurotransmitter acetylcholine (ACh) to terminate ACh action at the NMJ. The aggregation and precise alignment of AChE with AChRs at post-synaptic membrane is vital for neurotransmission. We double-labeled wholemounts of Sol, EDL, and TS muscles with antibodies against AChE and Texas Red-conjugated *α*-bungarotoxin. We found that AChE staining was completely colocalized with the AChR staining in both *WT* and *Acta1*^+*/Ki*^ muscles (Fig. 2).

**Figure 2.**
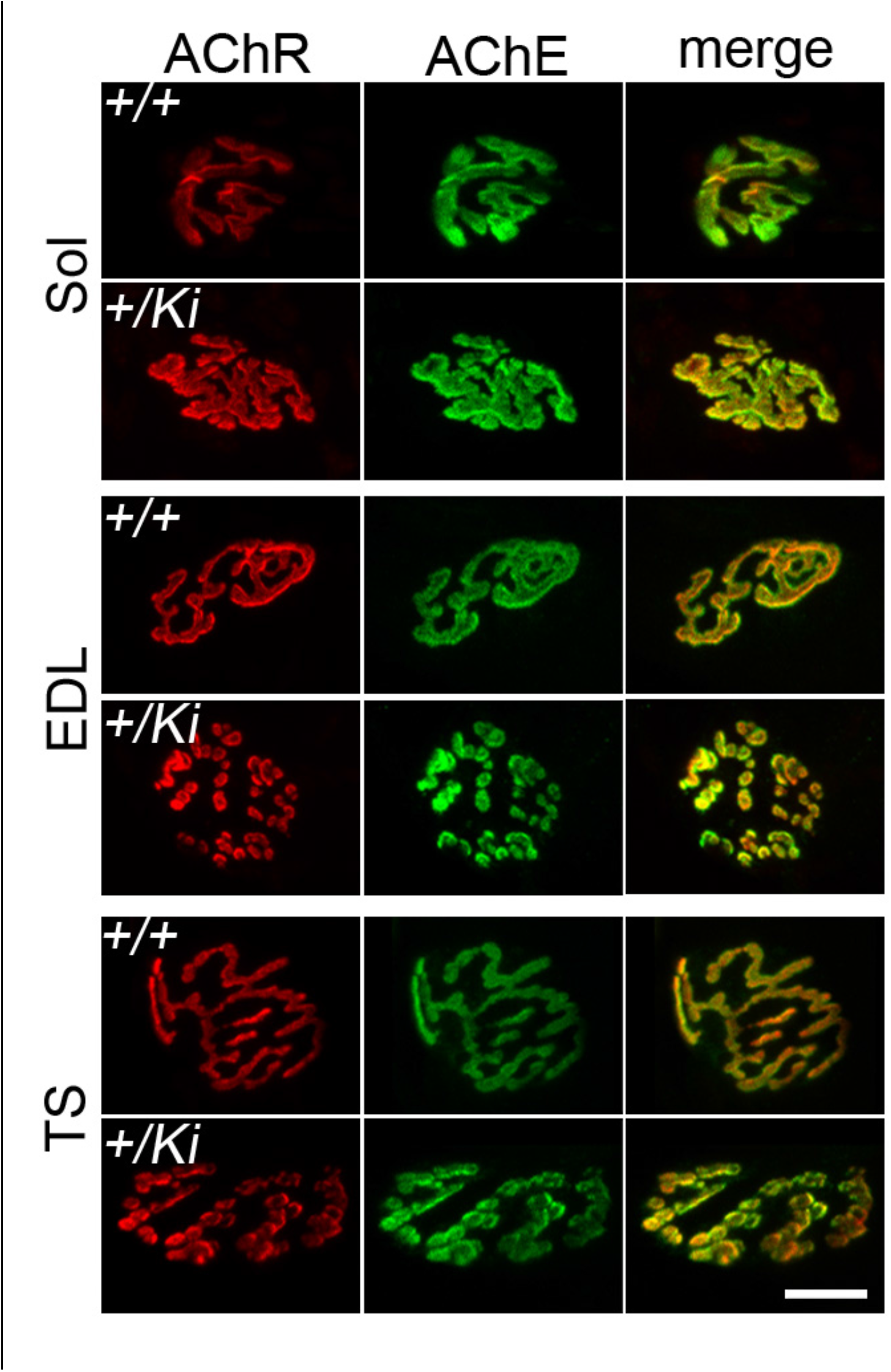
Fragmentation of AChE clusters *in Acta1*^+*/Ki*^ mice. Whole mounts of Sol, EDL and TS muscles of *Acta1*^+*/Ki*^ and *WT* mice (2-month-old) were double-labeled with Texas Red-conjugated *α*-bungarotoxin for AChRs (red) and antibodies against AChE (green). Fragmentation was detected In *Acta1*^+*/Ki*^ EDL and TS muscles. Scale bars: 20 μm.

### Presynaptic nerves are altered at the NMJs of *Acta1*^+/Ki^ mice

We next examined presynaptic nerve terminals using antibodies against synaptic vesicle proteins such as anti-synaptotagmin 2 (syt 2). Consistent with the alterations seen at postsynaptic AChRs, pre-synaptic nerve terminals in *Acta1*^+*/Ki*^ mice appeared fragmented and exhibited a bead-like staining pattern in EDL and TS muscles (arrowheads in Fig. 3B-C), but not in Sol muscles (Fig. 3A). This was quantified as the ratio of nerve occupancy by dividing the area of the nerve terminal by the area occupied by AChRs. The nerve occupancy ratio was comparable in Sol muscles of *WT* and *Acta1*^+*/Ki*^ mice. However, the nerve occupancy ratio in *Acta1*^+*/Ki*^ mice was significantly reduced in EDL and TS muscles compared to in *WT* (Fig. 3E). Nevertheless, in *Acta1*^+*/Ki*^ muscles presynaptic nerve terminals were juxtaposed with the postsynaptic endplate; no denervated endplates were detected.

**Figure 3.**
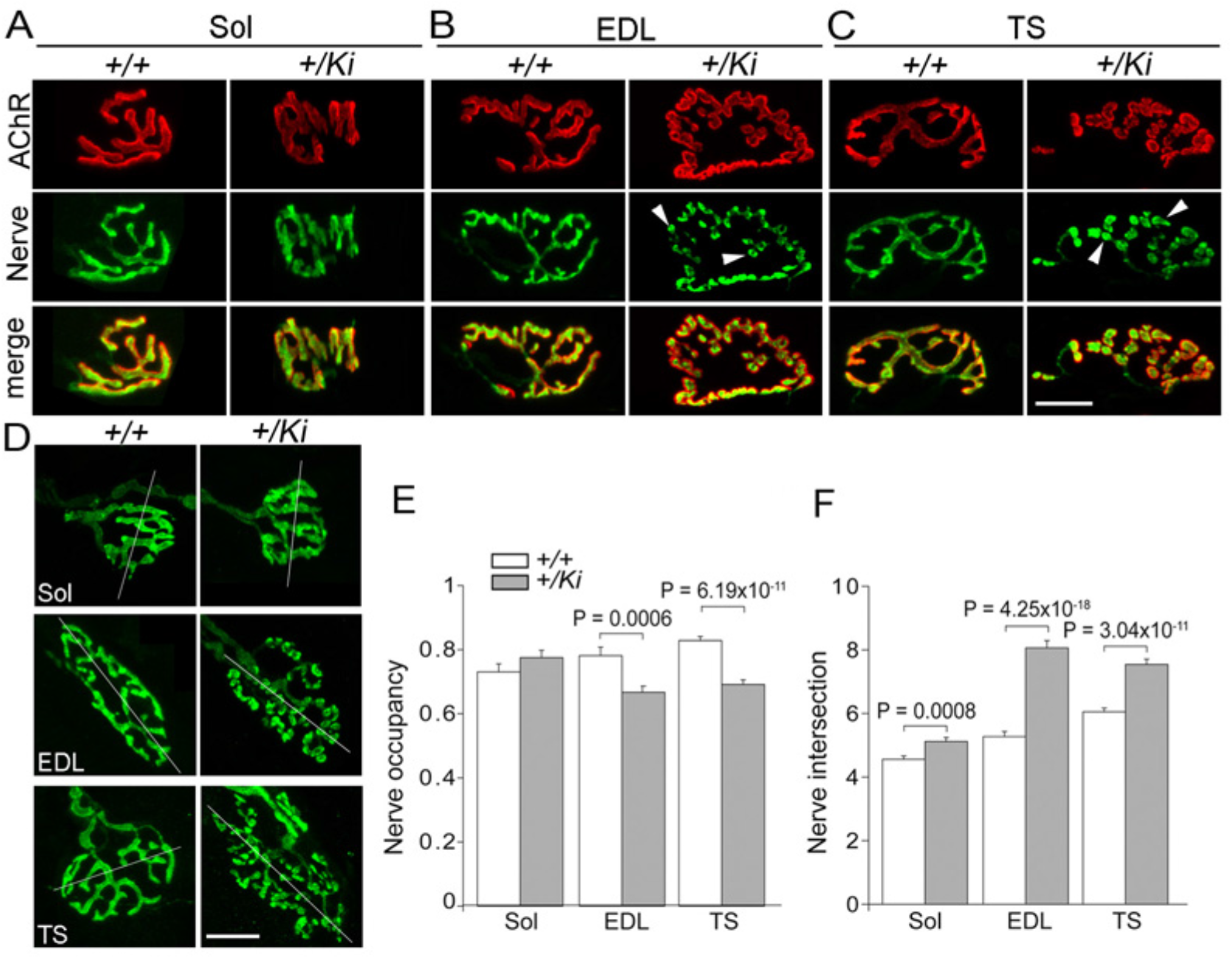
Pre-synaptic abnormalities *in Acta1*^+*/Ki*^ mice. A-C: whole mounts of Sol (A), EDL (B) and TS (C) muscles of *Acta1*^+*/Ki*^ and *WT* mice (2-month-old) were double labeled with antibodies against synaptotagmin 2 for nerve terminals and Texas Red-conjugated *α*-bungarotoxin for AChR clusters. D: whole mounts of Sol, EDL and TS muscles were labeled with antibodies against syntaxin 1 for preterminal axon. A line was drawn along the longest axes of the neuromuscular junction and the number of the intersections between the line and axons was counted. E: Nerve occupancy was calculated as a ratio of nerve terminal area over AChR area. Nerve occupancy in the Sol muscle was normal in *Acta1*^+*/Ki*^ mice (0.78 ± 0.02) compared to *WT* (0.73 ± 0.03). In contrast, nerve occupancy was significantly reduced in EDL (*Acta1*^+*/Ki*^ 0.67 ± 0.02; WT: 0.78 ± 0.03) and TS (*Acta1*^+*/Ki*^ 0.69 ± 0.01; WT: 0.83 ± 0.01). F: Average numbers of the nerve intersections obtained from D. No significant difference was detected in Sol between *Acta1*^+*/Ki*^ and *WT* mice. In contrast, the number of nerve intersection was notably increased in EDL (*Acta1*^+*/Ki*^ 8.06 ± 0.23 vs WT: 5.27 ± 0.16) and TS (*Acta1*^+*/Ki*^ 7.54 ± 0.17 vs WT: 6.05 ± 0.12). The numbers of animal (N) and neuromuscular junction (n) used for quantitative analysis: Sol (*WT*: N = 3, n = 134; *Acta1*^+*/Ki*^: N = 3, n = 119). EDL (*WT*: N = 3, n = 84; *Acta1*^+*/Ki*^: N = 3, n = 109). TS (*WT*: N = 3, n = 115; *Acta1*^+*/Ki*^: N = 3, n = 137). Scale bars: A-D, 20 μm.

Using anti-syntaxin1 antibodies to label pre-terminal axons, we noticed that pre-terminal axons appeared more complex in *Acta1*^+*/Ki*^ mice when compared to *WT* (Fig. 3D). To quantify the complexity, we drew a line along the longest axis across the nerve terminal and counted the number of intersections between the line and the axon labeled by the syntaxin antibody (Fig. 3D). The number of nerve intersects was markedly increased in *Acta1*^+*/Ki*^ mice (increase by 12% in Sol, 53% in EDL, and 25% in TS respectively, P<0.005, student *t*-test), compared to in *WT* (Fig. 3F). Such an increase in nerve branching has previously been reported in aging (Jang & Van Remmen, 2011; Li *et al*., 2018) and dystrophic mice (such as Duchenne muscular dystrophy) (Pratt *et al*., 2015), both of which also exhibit endplate fragmentation.

### The number of subsynaptic nuclei is increased in *Acta*1^+*/Ki*^ muscles

Nuclei within the synaptic region of a muscle fiber (subsynaptic nuclei) contribute to subsynaptic gene expression and therefore are transcriptionally distinct from those in the extra-synaptic region (Schaeffer *et al*., 2001; Hippenmeyer *et al*., 2007). Subsynaptic nuclei are defined as those within the synaptic region or that cross the boundary of the synaptic region (Pratt *et al*., 2015). We counted the numbers of subsynaptic nuclei and normalized this number to the area of the synaptic region (nuclei number per 100 μm^2^ synaptic area). We found that the density of subsynaptic nuclei was significantly increased in *Acta1*^+*/Ki*^ mice when compared with *WT* (increase by 46% in Sol, 51% in EDL, and 25% in TS respectively, P<0.005, student *t*-test) (Fig. 4C).

**Figure 4.**
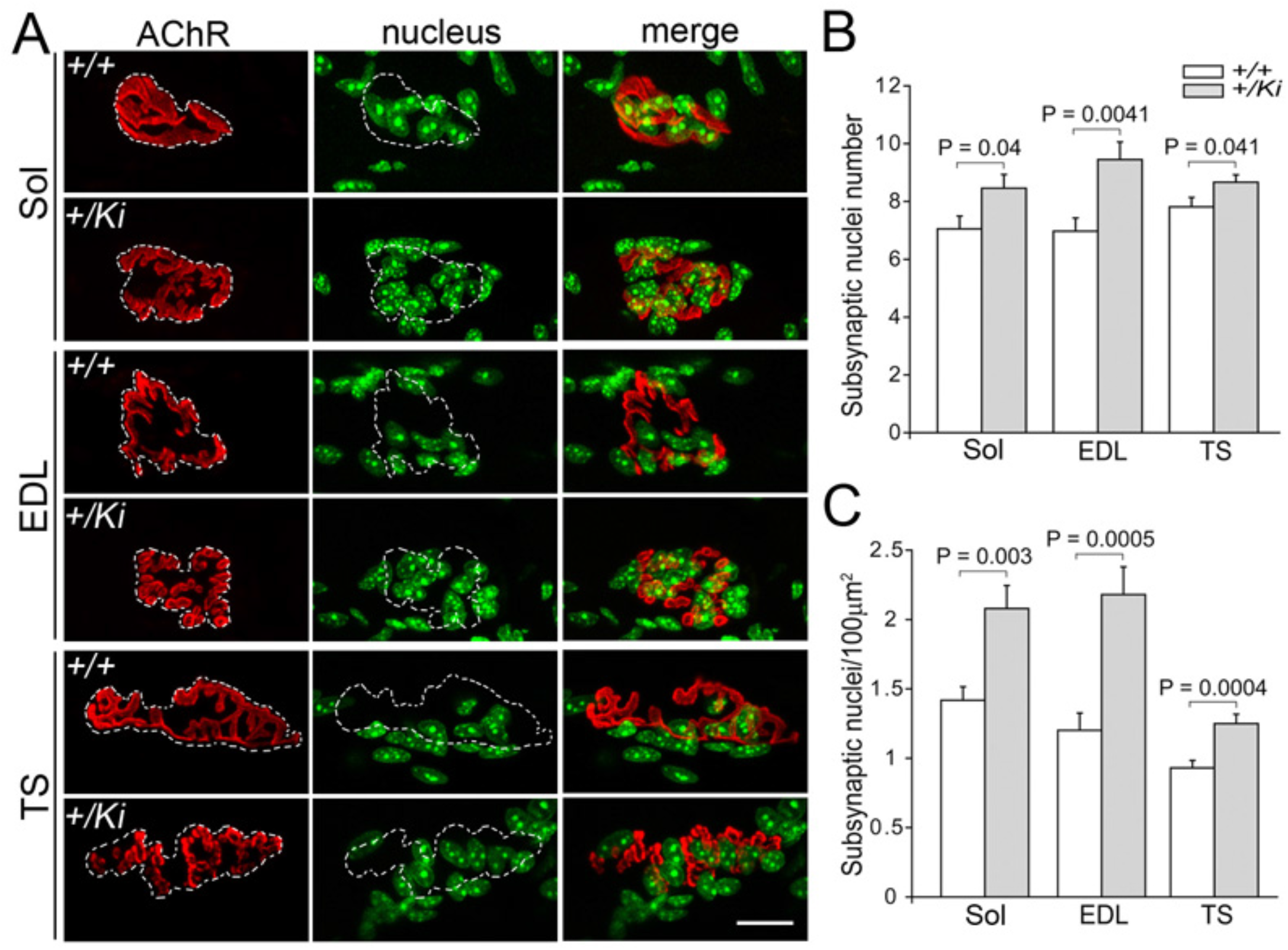
Increased subsynaptic nuclei in *Acta1*^+*/Ki*^ mice. A: whole mounts of Sol, EDL and TS muscles of *Acta1*^+*/Ki*^ and *WT* mice (2-month-old) were double-labeled with Texas Red-conjugated *α*-bungarotoxin for endplates (red) and Topro-3 for nuclei (green). Dotted lines indicate endplate region. B: Numbers of subsynaptic nuclei in Sol, EDL and TS muscles. Compared to those of control muscles, the numbers of subsynaptic nuclei were significantly increased in three mutant muscles. (Sol, *Acta1*^+*/Ki*^: 8.46 ± 0.47 vs *WT* 7.05 ± 0.44; EDL, *Acta1*^+*/Ki*^: 9.45 ± 0.61 vs *WT* 6.97 ± 0.46; TS: *Acta1*^+*/Ki*^: 8.67 ± 0.25 vs *WT*: 7.81 ± 0.33). C: Normalized number of subsynaptic nuclei to endplate region (nuclei/100 μm^2^). Compared to those of control muscles, the normalized numbers of subsynaptic nuclei were significantly increased in all three muscles (Sol, *Acta1*^+*/Ki*^: 2.08 ± 0.17 vs *WT* 1.42 ± 0.1; EDL, *Acta1*^+*/Ki*^: 2.17 ± 0.19 vs *WT* 1.44 ± 0.15; TS: *Acta1*^+*/Ki*^: 1.16 ± 0.06 vs *WT*: 0.93 ± 0.05 Scale bar: 20 μm.

### Spontaneous and evoked transmitter release is changed at the NMJs of *Acta1*^+*/Ki*^ mice

We carried out intracellular recording on EDL and Sol muscles of *Acta1*^+*/Ki*^ and *WT* mice at 2 months of age. The resting membrane potentials were comparable between *WT* and *Acta1*^+*/Ki*^ muscles: *WT* (EDL: −70.2 ± 0.8 mV, N = 3 mice, n= 43 cells; Sol: −69.9 ± 1 mV, N = 3, n = 41); Acta1^+/*Ki*^ (EDL: −67.9 ± 0.8 mV, N = 3, n = 46; Sol: −67.6 ± 0.7 mV, N =3, n = 38). We next recorded miniature endplate potentials (mEPPs) at resting state, resulting from spontaneous neurotransmitter release of synaptic vesicles from unstimulated nerve terminals. In *Acta1*^+*/Ki*^ mice, the frequency of mEPP increased by 29% in EDL muscles (1.27 ± 0.09 Hz) and 40% in Sol muscles (1.92 ± 0.16 Hz), compared to *WT* (EDL: 0.98 ± 0.11 Hz; Sol: 1.36 ± 0.11 Hz), P<0.05 by student *t*-test. However, mEPP amplitudes were reduced by 23% in EDL muscles (0.63 ± 0.03 mV) and 15% in Sol muscles (0.84 ± 0.04 mV) in *Acta1*^+*/Ki*^ mice, compared to *WT* (EDL: 0.82 ± 0.05 mV; Sol: 0.98 ± 0.04 mV), P<0.05 by student *t*-test. The 10~90% rise time was significantly longer in both EDL muscles (1.73 ± 0.08 ms) and Sol muscles (2.02 ± 0.12 ms) in *Acta1*^+*/Ki*^ mice, compared to *WT* (EDL: 1.34 ± 0.12 ms; soleus: 1.57 ± 0.07 ms) (Fig. 5), P<0.05 by student *t*-test.

**Figure 5.**
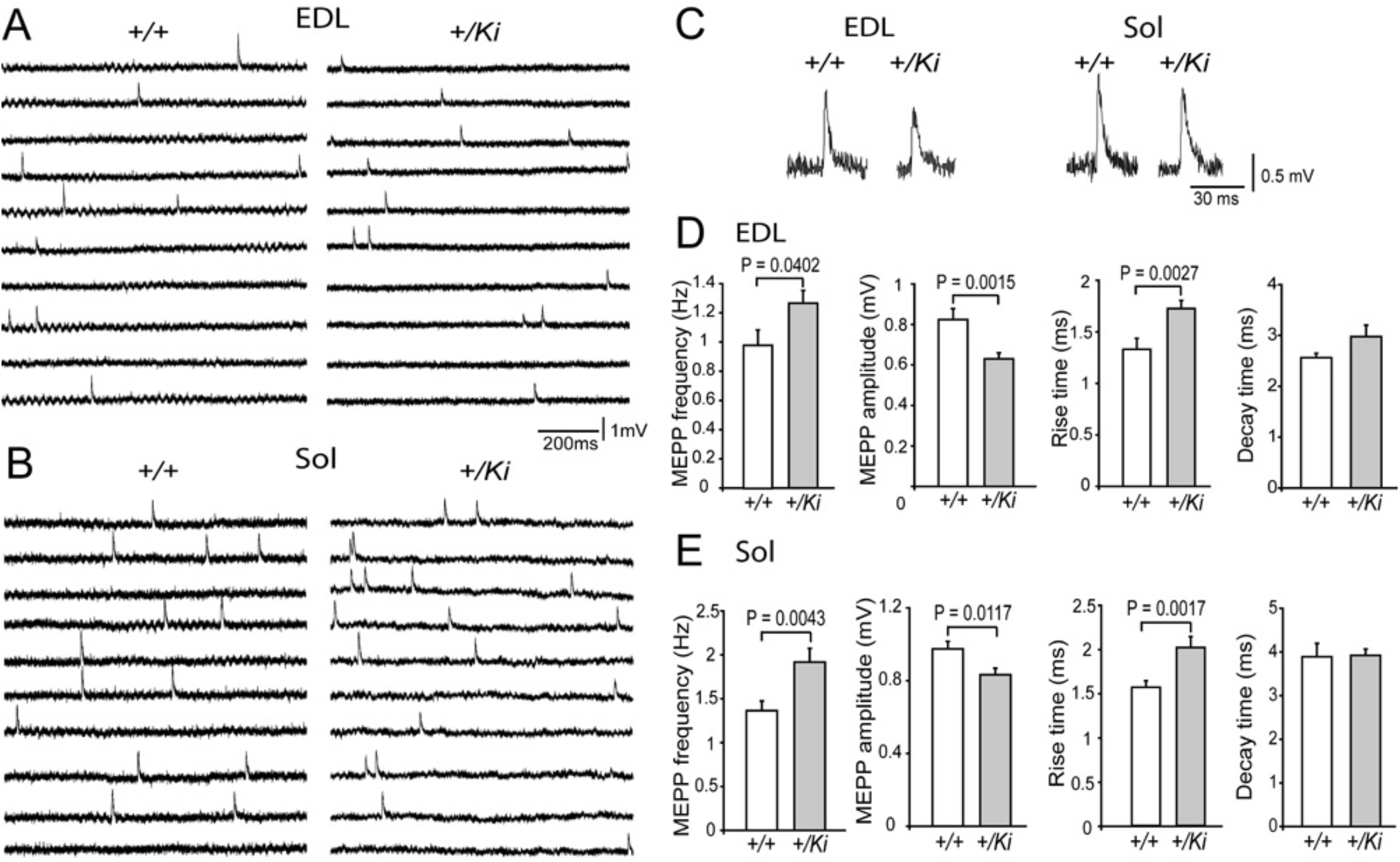
Alterations in spontaneous transmitter release at NMJs in *Acta1*^+*/Ki*^ mice. A, B: Sample mEPPs recorded in EDL (A) and Sol (B) of *Acta1*^+*/Ki*^ and *WT* mice (2-month-old). C: Individual mEPP traces recorded in EDL and Sol muscles. D: Quantification of mEPP frequency, amplitude, rise time (10%~90%) and decay time (100%~50%) in EDL. MEPP frequency was notably higher in *Acta1*^+*/Ki*^ mice (1.27 ± 0.09 Hz) than that in *WT* (0.98 ± 0.11 Hz). MEPP amplitude was significantly reduced in *Acta1*^+*/Ki*^ mice (0.63 ± 0.03 mV) compared with that in *WT* (0.82 ± 0.05 mV). MEPP rise time was markedly increased in *Acta1*^+*/Ki*^ mice (1.73 ± 0.08 ms) compared with that in *WT* (1.34 ± 0.12 ms). MEPP decay time was comparable between *Acta1*^+*/Ki*^ (2.98 ± 0.23 ms) and *WT* (2.56 ± 0.09 ms) mice. E: Quantification of mEPP frequency, amplitude, rise time (10%~90%) and decay time (100%~50%) in Sol. MEPP frequency was significantly increased in *Acta1*^+*/Ki*^ mice (1.92 ± 0.16 Hz) compared with that in control mice (1.36 ± 0.11 Hz). MEPP amplitude was significantly reduced in *Acta1*^+*/Ki*^ mice (0.84 ± 0.04 mV) compared with that in *WT* mice (0.98 ± 0.04 mV). MEPP rise time was notably increased in *Acta1*^+*/Ki*^ mice (2.02 ± 0.12 ms) compared with that in *WT* (1.57 ± 0.07 ms). MEPP decay time in *Acta1*^+*/Ki*^ (3.92 ± 0.15 ms) was similar with that in that in *WT* mice (3.89 ± 0.31 ms). The numbers of animal (N) and muscle fibers (n) used for analysis: EDL (*WT*: N = 3, n=43; *Acta1*^+*/Ki*^: N = 3, n = 46). Sol (*WT*: N = 3, n = 41; *Acta1*^+*/Ki*^: N = 3, n = 38).

Endplate potentials (EPPs) were recorded to evaluate neurotransmitter release evoked by nerve action potentials. Intriguingly, unlike the reduced amplitude in mEPPs that was observed previously, there was no significant change in EPP amplitude between *Acta1*^+*/Ki*^ and *WT* mice in both EDL and Sol muscles. However, the quantal content, which represents the quantal number of transmitter release in response to a nerve impulse, was increased by 36% in EDL muscles (32.43 ± 1.72) and 13% in Sol muscles (34.7 ± 1.41) in *Acta1*^+*/Ki*^ mice, compared to in *WT* mice (EDL: 23.8 ± 1.55; Sol: 30.68 ± 0.98), P<0.05, student *t*-test. We noted in both *WT* and mutant mice, the amplitude of EPPs recorded in Sol muscles was significantly larger than that of EDL muscles, consistent with previous reports (Banker *et al*., 1983).

### Short-term synaptic plasticity is impaired in *Acta1*^+*/Ki*^ mice

Short-term plasticity, either facilitation or depression, refers to the changes in synaptic strength over a short time scale (within hundreds to thousands of milliseconds). First, we examined short-term facilitation by applying paired pulses in variable intervals from 20-50 ms to the nerve and recorded the evoked EPPs in the muscle. The second EPP was larger than the first EPP due to the residual Ca^2+^ in the nerve terminal caused by the first nerve impulse (Zucker & Regehr, 2002; Jackman & Regehr, 2017). The amplitude ratio of the second EPP to the first EPP indicates the degree of facilitation. Compared to *WT*, *Acta1*^+*/Ki*^ EDL and Sol muscles exhibited smaller ratios of EPP(2)/EPP(1) (Fig. 7C-D).

**Figure 6.**
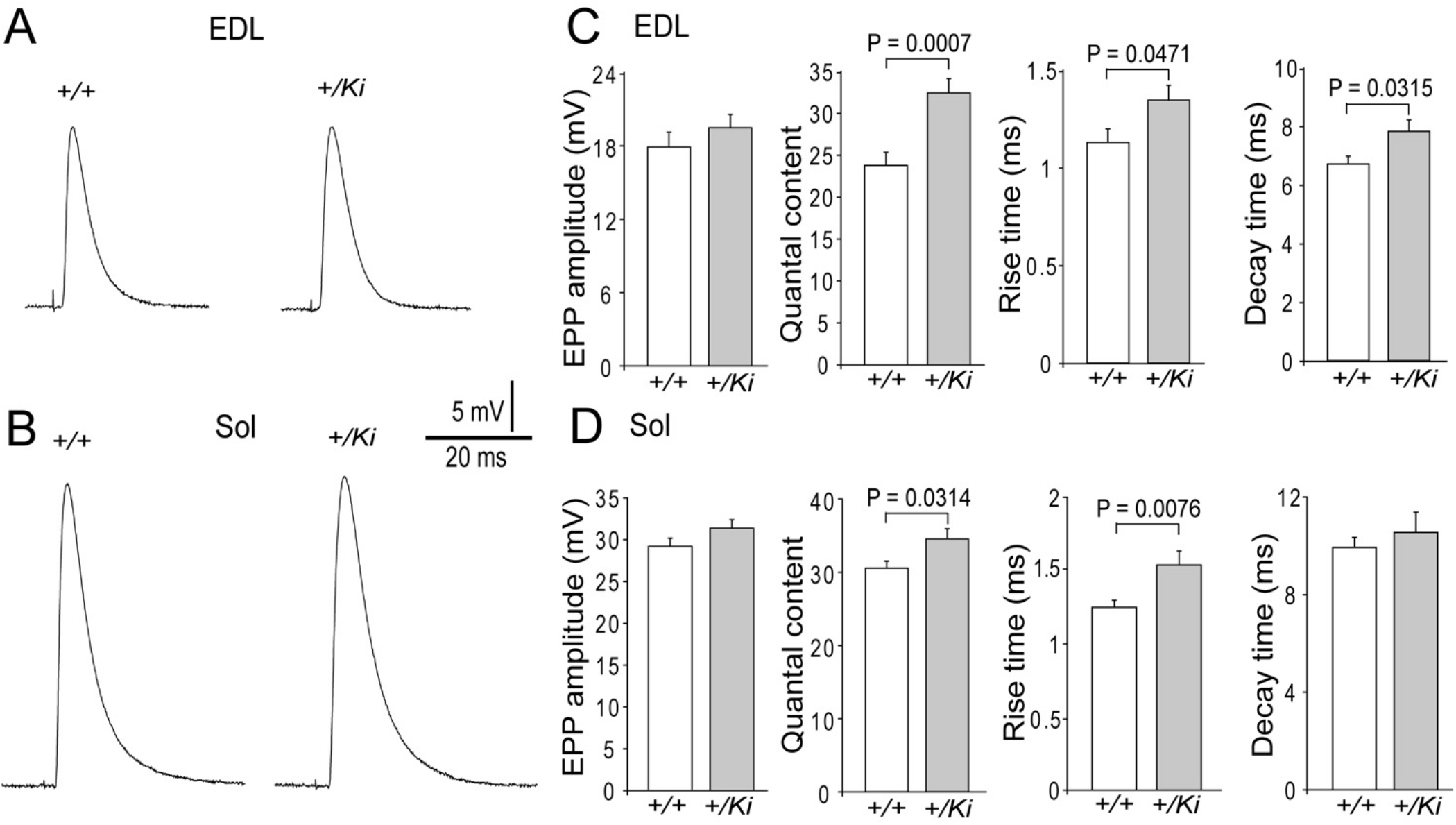
Alterations in evoked transmitter release at NMJs in *Acta1*^+*/Ki*^ mice. A, B: Sample EPPs recorded in EDL (A) and Sol (B) muscles of *Acta1*^+*/Ki*^ and *WT* mice (2-month-old). C: Quantification of EPP amplitude, quantal content, rise time (10%~90%) and decay time (90%~10%) in EDL. EPP amplitude was comparable in *Acta1*^+*/Ki*^ mice (19.49 ± 1.08 mV) and *WT* (17.9 ± 1.21 mV). Quantal content was significantly increased in *Acta1*^+*/Ki*^ mice (32.43 ± 1.72) compared with that in *WT* (23.8 ± 1.55). Rise time and decay time were significantly increased in *Acta1*^+*/Ki*^ mice (1.35 ± 0.08 ms and 7.87 ± 0.39 ms, respectively) compared with those in *WT* (1.13 ± 0.07 ms and 6.75 ± 0.27 ms, respectively). D: Quantification of EPP amplitude, quantal content, rise time (10%~90%) and decay time (90%~10%) in Sol. EPP amplitude was similar between *Acta1*^+*/Ki*^ mice (31.38 ± 1.02 mV) and *WT* (29.21 ± 0.98 mV). Quantal content was notably increased in *Acta1*^+*/Ki*^ mice (34.7 ± 1.41) compared with that in *WT* (30.68 ± 0.98). EPP rise time was significantly increased in *Acta1*^+*/Ki*^ mice (1.54 ± 0.1 ms) compared with that in *WT* (1.25 ± 0.05 ms). EPP decay time was similar between *Acta1*^+*/Ki*^ (10.55 ± 0.84 mV) and *WT* (9.93 ± 0.41 mV). The numbers of animal (N) and muscle fibers (n) used for analysis: EDL (*WT*: N = 3, n = 24; *Acta1*^+*/Ki*^: N = 3, n = 32); Sol (*WT*: N = 3, n = 24; *Acta1*^+*/Ki*^: N = 3, n = 23).

**Figure 7.**
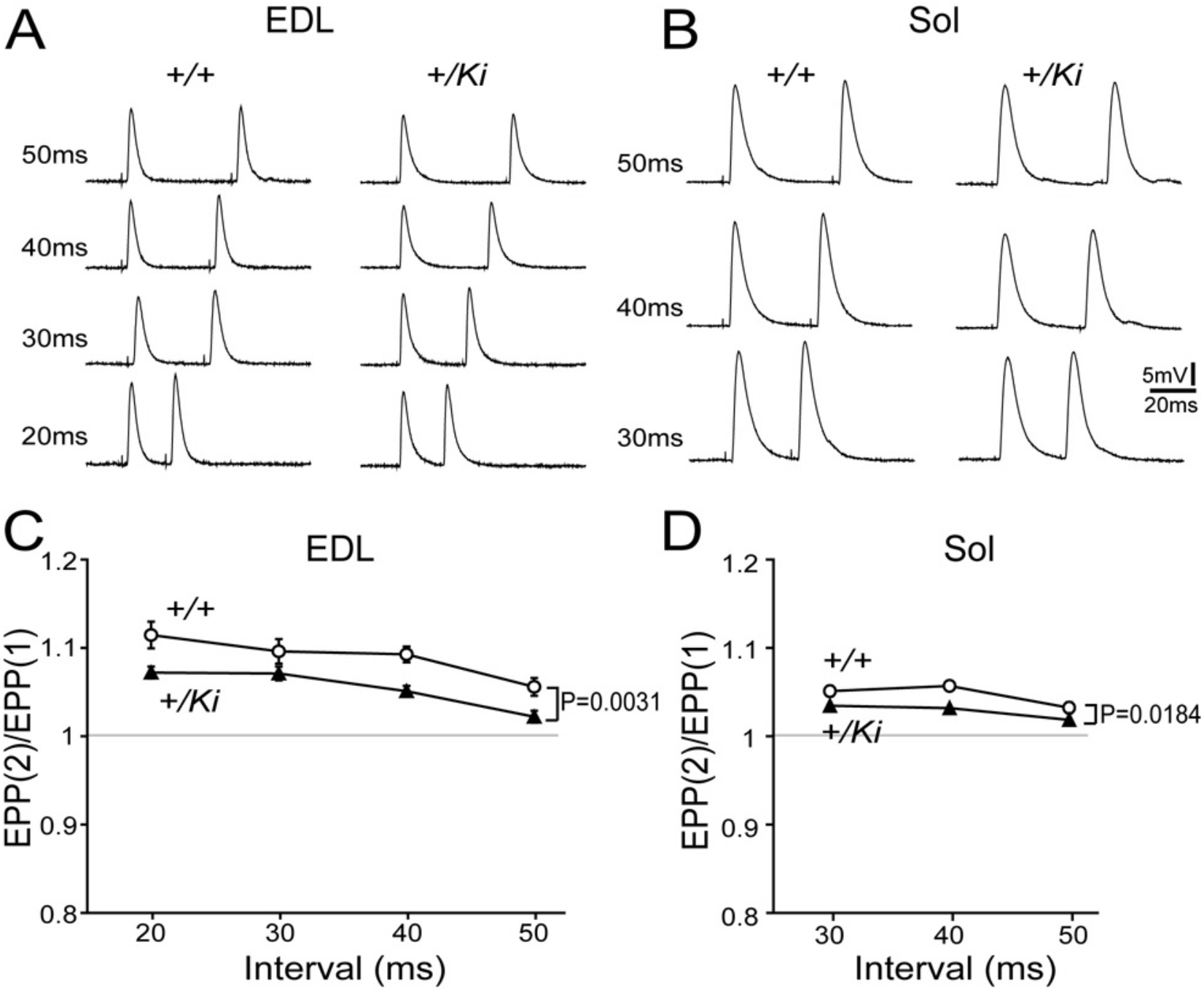
Reduced paired-pulse facilitation in *Acta1*^+*/Ki*^ mice. A, B: Sample EPPs recorded in EDL (A) and Sol (B) of *Acta1*^+*/Ki*^ and *WT* mice (2-month-old) in response to pair-pulse stimulation to the nerve at various interpulse intervals (20-50ms). C-D: The ratios of the second to the first EPP amplitude (EPP(2)/EPP(1)). Pair-pulse facilitation was significantly reduced at all intervals in both EDL and Sol in *Acta1*^+*/Ki*^ mice. The numbers of animal (N) and muscle fibers (n) used for analysis: EDL (*WT*: N = 3, n = 22; *Acta1*^+*/Ki*^: N = 3, n = 32). Sol (*WT*: N = 3, n = 22; *Acta1*^+*/Ki*^: N = 3, n = 27).

Synaptic depression occurs when synaptic vesicles in the nerve terminal are depleted during repetitive nerve activity (Zucker & Regehr, 2002). To determine whether such depression was altered in mutant mice, we applied high frequency stimulation (30 Hz) to the nerve and recorded EPPs in the muscle. In *WT* muscles, EPPs initially exhibited moderate facilitation, then rapidly progressed to depression before eventually reaching a plateau. EPPs recorded in *Acta1*^+*/Ki*^ EDL and Sol muscles showed patterns like those in *WT* muscles but exhibited significantly greater depression before reaching a plateau (Fig. 8C-D). These results indicate that short-term synaptic plasticity was compromised in *Acta1*^+*/Ki*^ mice.

**Figure 8.**
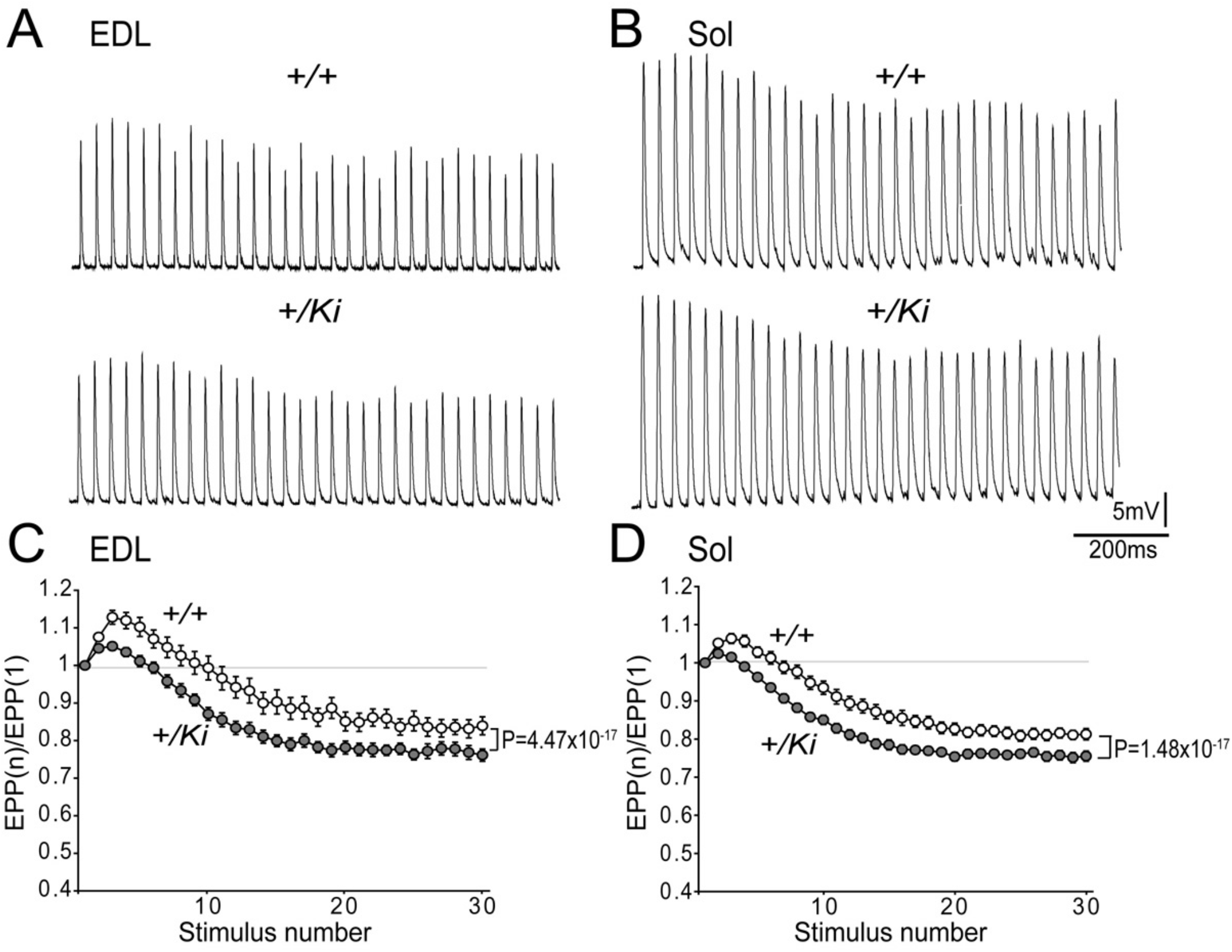
Increased synaptic depression in *Acta1*^+*/Ki*^ mice. A-B: Sample EPPs recorded in EDL (A) and Sol (B) of *Acta1*^+*/Ki*^ and *WT* mice (2-month-old) in response to 1-second of 30 Hz train stimulation of the nerve. C-D: Both *Acta1*^+*/Ki*^ EDL and Sol exhibited significantly increased depression following train stimulation, compared with those of control muscles. The numbers of animal (N) and muscle fibers (n) used for analysis: EDL (*WT*: N = 3, n=21; *Acta1*^+*/Ki*^: N = 3, n = 27). Sol (*WT*: N = 3, n = 24; *Acta1*^+*/Ki*^: N = 3, n = 28).

## Discussion

In this study, we report changes in the NMJs of *Acta1*^+*/Ki*^ mice, which exhibit clinical features of human congenital myopathy, including premature lethality, severe muscle weakness, reduced mobility, and the presence of nemaline rods in muscle fibers (Nowak *et al*., 1999; Nguyen *et al*., 2011). We found that the development and formation of the NMJ were initially normal. However, maintenance of the NMJ was impaired in the *Acta1*^+*/Ki*^ mice. The alterations included endplate fragmentation, reduced or loss of AChRs, increased nerve terminal complexity, decreased nerve occupancy, and increased number of subsynaptic nuclei. Furthermore, our electrophysiological analyses revealed reduced mEPP amplitude but normal EPP size and altered short-term synaptic plasticity at the NMJs of *Acta1*^+*/Ki*^ mutant mice.

During postnatal development of the NMJ, the AChR cluster is gradually sculpted from an oval “plaque” into curved bands with a “pretzel” shape (Marques *et al*., 2000). Correspondingly, the presynaptic nerve terminal is elaborated to juxtapose against the AChR clusters. The complex structure ensures reliable neurotransmission at the NMJ. Our data show that the continuous, “pretzel” structure of the endplate become fragmented into many discrete islets in *Acta1*^+*/Ki*^ mice as early as one month of age. The causes of NMJ fragmentation remain unclear. But it has also been observed in aged and dystrophic muscle (Harris & Ribchester, 1979; Prakash & Sieck, 1998; Gonzalez-Freire *et al*., 2014; Rudolf *et al*., 2014; Pratt *et al*., 2015; Lovering *et al*., 2020). Several lines of evidence suggest that NMJ fragmentation could be due to synaptic remodeling caused by the regeneration of deteriorated muscle fiber segments underlying the affected synapse (Li *et al*., 2011; Li & Thompson, 2011). Additionally, it has also been suggested that NMJ fragmentation is a feature of regeneration, and therefore a normal process by which the efficacy of the NMJ is maintained during aging or diseases (Slater, 2020). The mutation H40Y in α-actin disrupts the binding between actin in the thin filament and myosin in the thick filament, which causes reduced muscle force and leads to severe muscle weakness (Chan *et al*., 2016). Signs of focal myofiber damage (lack of eosin staining in some myofiber areas) and chronic repair and regeneration (muscle fibers with internal nuclei) have been observed in *Acta1*^+*/Ki*^ mice (Nguyen *et al*., 2011). In addition, our data show that the number of nuclei within muscle subsynaptic regions is markedly increased, suggesting that regeneration and remodeling may be occurring at the synaptic site in *Acta1*^+*/Ki*^ mice. Although the direct effect of the *Acta1* (H40Y) mutation on the central nervous system remains unknown, no loss of neurons has been found in patient with nemaline myopathy caused by *ACTA1* mutation (Asp154Asn) (Schroder *et al*., 2004). Thus, it is plausible that the endplate fragmentation in *Acta1*^+*/Ki*^ mice is attributable to myofiber degeneration and regeneration, and that presynaptic nerve alterations could be secondary to postsynaptic disruptions at the NMJ.

We found that the severity of NMJ fragmentation varied greatly among different muscle types. Endplate fragmentation was most severe in EDL, relatively moderate in TS, and least affected in Sol. The difference in susceptibility to NMJ fragmentation could be due to the different extent of myopathies in different muscle fiber types in *Acta1*^+*/Ki*^ mutant mice. The EDL is a fast twitch muscle which is predominantly composed of Type IIb fibers, whereas the Sol is a slow twitch muscle mainly consisting of Type I and IIa fibers (Schiaffino & Reggiani, 2011). Indeed, a previous study has reported different muscles in *Acta1*^+*/Ki*^ mice exhibit different extents of muscle damage (Nguyen *et al*., 2011). While muscle fibers with internal nuclei are detected in both EDL and Sol muscles *Acta1*^+*/Ki*^ mice, the percentage of fibers with internal nuclei is significantly higher in EDL than in *Sol*. In addition, as a common feature of nemaline myopathy, the shift towards slow fiber types (an increase in type I fiber and a concomitant decrease in type IIa fibers) is also observed in the Sol muscle of *Acta1*^+*/Ki*^ mice (Nguyen *et al*., 2011). Analogous muscle and fiber-type specificity in NMJ morphological alteration has been observed during aging and in neuromuscular diseases (Prakash & Sieck, 1998; Valdez *et al*., 2012). For example, NMJs in EDL muscle are highly susceptible to aging; however, NMJs in extraocular muscle are strikingly resistant to damages (Valdez *et al*., 2012). Similarly, fast muscle fibers (type IIb) have been shown to degenerate first in Duchenne muscular dystrophy (Webster *et al*., 1988).

A disrupted NMJ structure may undermine the efficacy of neuromuscular synaptic transmission. The alterations in neurotransmission that we observed in the *Acta1*^+*/Ki*^ mice, such as reduced mEPP amplitude (quantal size) and compromised short-term plasticity, suggest functional impairments. At postsynaptic sites, mEPP size is positively correlated to postsynaptic AChR density and negatively correlated to muscle fiber size (Katz & Thesleff, 1957; Harris & Ribchester, 1979). As muscle fiber sizes are significantly reduced in *Acta1*^+*/Ki*^ mice (Nguyen *et al*., 2011), an increase in mEPP amplitude is expected based on the correlation of fiber size and mEPP amplitude. However, we found that the mEPP amplitude was significantly reduced in *Acta1*^+*/Ki*^ mice. Thus, the decrease in mEPP size likely resulted from a reduction or loss of ACh receptors on the muscle membrane, as demonstrated by our morphological analyses.

Intriguingly, mEPP frequencies were significantly increased in *Acta1*^+*/Ki*^ mice. The elevated mEPP frequency suggests that compensatory adjustments occurred at affected synapses to increase presynaptic transmitter release in order to compensate for reduced postsynaptic excitation caused by a reduction in AChRs (Davis, 2013; Davis & Muller, 2015). This presynaptic homeostasis is considered to be crucial to restore synapse baseline function in the presence of perturbations. Furthermore, *Acta1*^+*/Ki*^ mutant NMJs exhibited increased quantal content and thus normal EPP amplitude, which further indicated that synaptic homeostatic modulations occurred at *Acta1*^+*/Ki*^ mutant synapses to enhance the transmitter release per nerve impulse in order to offset the reduction in AChRs and maintain the safety factor of the NMJ (Ruff, 2011; Davis & Muller, 2015). Synaptic homeostatic compensatory mechanisms are involved in aging (Banker *et al*., 1983; Willadt *et al*., 2016) as well as in various neuromuscular diseases such as myasthenia gravis (MG) (Plomp *et al*., 2015) and Duchenne muscular dystrophies (van der Pijl *et al*., 2016). The molecular mechanisms underlying presynaptic homeostasis remain to be further elucidated. It has been suggested that calcium influx signals in presynaptic nerve terminals and release-ready vesicle pools are involved (Muller & Davis, 2012; Muller *et al*., 2012; Davis & Muller, 2015).

The endplate fragmentation in *Acta1*^+*/Ki*^ mice appears to be muscle fiber type dependent, that is, fasttwitch muscles (EDL) are more susceptible whereas slow-twitch muscles (Sol) are less susceptible. In contrast, changes in NMJ function affected both fast (EDL) and slow (Sol) fiber muscles, and therefore appear to be independent of muscle group or fiber-type. Thus, endplate fragmentation in *Acta1*^+*/Ki*^ mice is not a direct indication of changes in NMJ function. This finding is of particular interest since it suggests that endplate fragmentation, which is considered to be a sign of synaptic degeneration and regeneration during aging (Jang & Van Remmen, 2011; Li *et al*., 2011) and neuromuscular diseases (Rudolf *et al*., 2014; Lovering *et al*., 2020), is not directly associated with impaired neurotransmission. Our results are in line with findings from a previous study demonstrating that NMJs in old mice (26-28 months-old), although substantially fragmented, do not exhibit a significant decline in neurotransmission compared with NMJs in middle-aged mice (12-14 month-old) (Willadt *et al*., 2016). Furthermore, increased quantal number per nerve impulse has been reported in EDL and Sol muscles in aged animals (Banker *et al*., 1983). Given the compensation of neurotransmitter release we observed in both mutant EDL and Sol muscles, it is plausible that the fragmented NMJs could restore and maintain the efficacy of neurotransmission through compensatory synaptic homeostatic mechanisms.

## Abbreviations

AChE: acetylcholinesterase
AChR: acetylcholine receptor
ACTA1: α_skeletal_-actin
EDL: extensor digitorum longus
EPP: endplate potential
MEPP: miniature endplate potential
NMJ: neuromuscular junction
Sol: soleus
TS: triangularis sterni

## Acknowledgments

We would like to thank Ms Qiaohong Ye for her excellent technical assistant, and Drs. Joseph McArdle, Beverly Rothermel, Jane Johnson and Ben Szaro for their critical comments on earlier drafts of the manuscript. This work was supported by grants from The Paul D. Wellstone Muscular Dystrophy Cooperative Research Center (HD-087351) (Y. Liu), NIH/NINDS (R01 NS055028) (W. Lin.).

